# Thermal niche evolution across replicated *Anolis* lizard adaptive radiations

**DOI:** 10.1101/069294

**Authors:** Alex R. Gunderson, D. Luke Mahler, Manuel Leal

## Abstract

Elucidating how ecological and evolutionary mechanisms interact to produce and maintain biodiversity is a fundamental problem in evolutionary ecology. We investigate this issue by focusing on how physiological evolution affects performance and species coexistence along the thermal niche axis in replicated radiations of *Anolis* lizards, groups best known for resource partitioning based on morphological divergence. We find repeated divergence in thermal physiology within these radiations, and that this divergence significantly affects performance within natural thermal environments. Morphologically similar species that co-occur invariably differ in their thermal physiology, providing evidence that physiological divergence facilitates species co-existence within anole communities. Despite repeated divergence in traits of demonstrable ecological importance, phylogenetic comparative analyses indicate that physiological traits have evolved more slowly than key morphological traits related to the structural niche. Phylogenetic analyses also reveal that physiological divergence is correlated with divergence in broad-scale habitat climatic features commonly used to estimate thermal niche evolution, but that the latter incompletely predicts variation in the former. We provide comprehensive evidence for repeated adaptive evolution of physiological divergence within *Anolis* adaptive radiations, including the complementary roles of physiological and morphological divergence in promoting community-level diversity. We recommend greater integration of performance-based traits into analyses of climatic niche evolution, as they facilitate a more complete understanding of the phenotypic and ecological consequences of climatic divergence.

## Introduction

A mechanistic understanding of how divergent phenotypic evolution facilitates species coexistence is a fundamental problem in ecology and evolution [1]. This problem comes into particular focus in the study of adaptive radiation, a process for which a hallmark is the repeated evolution of niche differences that allow closely related species to partition resources and stably co-occur [2-4]. Traditionally, most research on trait evolution during adaptive radiation has focused on morphological traits associated with dietary resource acquisition or structural habitat use, due to the clear role such adaptations play in mediating competition and facilitating coexistence [3, 5-7]. Physiological adaptation to abiotic conditions has received comparatively less attention in adaptive radiation studies, particularly in animals [3, 8]. Such adaptations are more typically studied for their role in generating divergence among geographically disjunct lineages experiencing different climates [9, 10]. However, physiological divergence can also play an important role in adaptive radiation by facilitating fine-scale partitioning of the local climatic niche [11], providing an additional resource axis along which a radiating clade may diversify [12, 13]. Indeed, such abiotic niche divergence has been hypothesized to underlie many impressive radiations, including bolitoglossine salamanders [14, 15], *Liolaemus* lizards [16-18], *Drosophila* flies [19], *Petrolisthes* crabs [20] and lake whitefish [21].

If physiological adaptation to abiotic conditions contributes to adaptive radiation, physiological divergence should affect performance in the different abiotic regimes that species occupy. Therefore, knowledge of both physiological traits related to the fundamental abiotic niche (broadly, the range of conditions a species could occupy) and abiotic conditions (at scales appropriate for the organisms under study [22]) are required to make the important phenotype-to-performance linkages established for many classic adaptive radiations driven by morphological divergence [3, 5, 23]. Despite the power of performance-based analyses, one of most common tools applied in recent investigations of abiotic niche divergence is correlative ecological niche modeling (ENM), which estimates species’ realized niches (*i.e.*, the range of conditions that species occupy) using information about species occurrences and broad-scale climate conditions [24-28]. Though correlative ENMs are powerful for addressing many questions, they also have characteristics that limit the inferences that can be made from their application [29-34]. In particular, correlative ENMs provide no information about phenotypic traits (physiological or otherwise) linked to performance [30]. Additionally, ENMs are limited in their ability to elucidate the role of abiotic niche divergence in coexistence, as they do not provide fine-scale information about microclimate use. Such factors can be of paramount importance for coexistence in radiating clades, as species with identical or largely overlapping ranges may be able to partition resources via adaptation to different microenvironments [11, 35].

Here, we investigate the contribution of thermal niche divergence to the replicated *Anolis* lizard adaptive radiations of Puerto Rico and Jamaica by analyzing patterns and performance consequences of physiological divergence across 16 species (Fig. 1). Greater Antillean *Anolis* are best known for morphological diversification along the fundamental structural niche axis [36-38], but morphological evolution cannot fully explain the patterns of divergence and community composition observed across the Greater Antilles [23]. Divergence in the thermal niche is also proposed to be an important component of *Anolis* diversification based primarily on realized niche estimates from natural history and ENM studies [12, 13, 24, 39-44]. However, we still know little about how physiology evolves during realized niche divergence [45] and the extent to which physiological evolution facilitates species co-existence.

**Figure 1:**
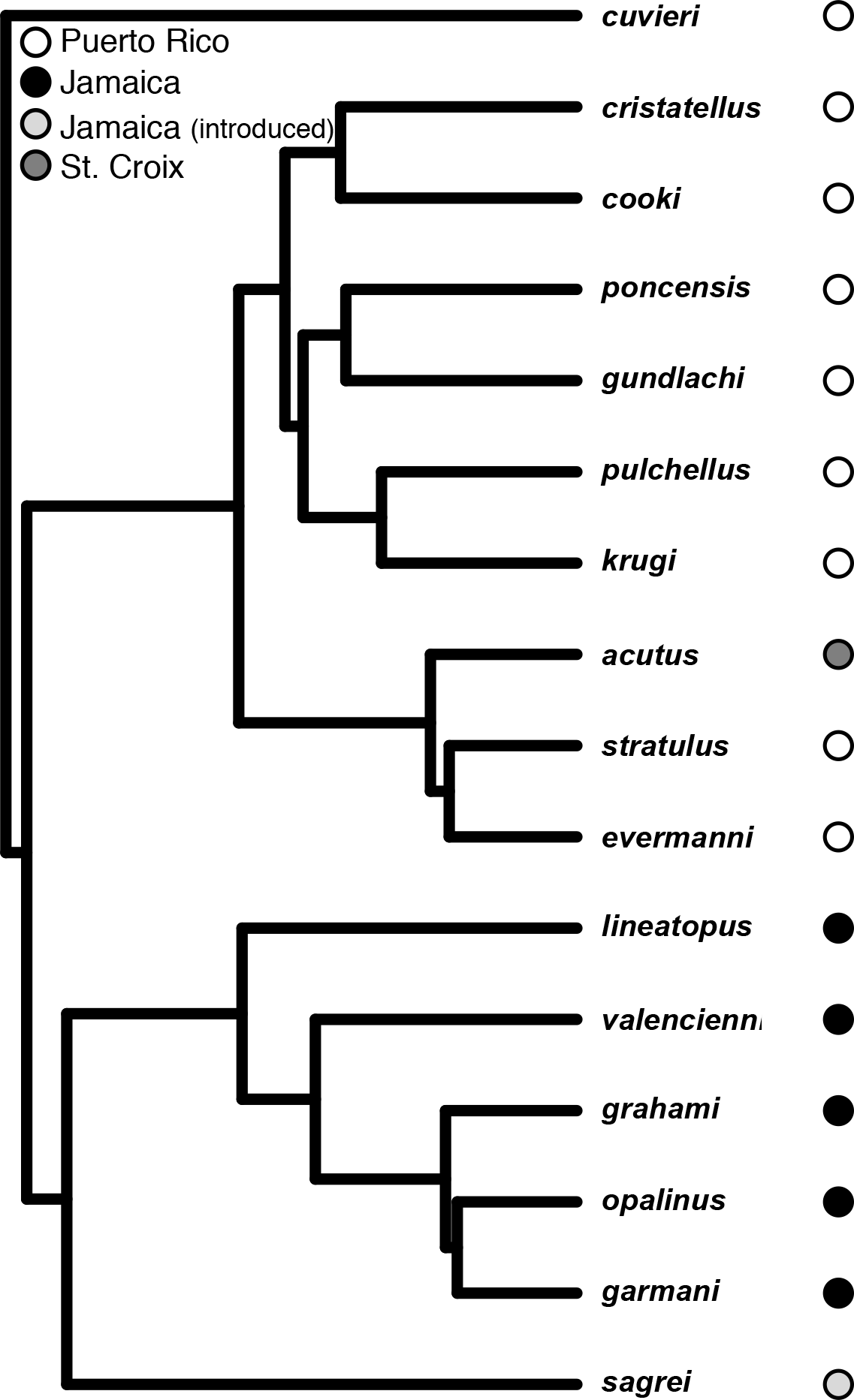
Phylogenetic relationships of the species included in this study, along with the islands from which they were sampled.

We first use data from the Puerto Rican *cristatellus* species group to investigate repeated divergence in thermal physiology and performance between species that occupy distinct thermal environments. Next, we integrate these data with measurements of operative thermal conditions in Puerto Rico to explore the performance consequences of physiological divergence. We then consider spatial patterns of physiological variation to evaluate the contribution of physiological evolution to fine-scale niche partitioning and community-level species co-existence across Puerto Rico and Jamaica. Finally, we conduct phylogenetic analyses of rates of physiological and morphological evolution to compare the pace of disparification between traits associated with the fundamental thermal niche and the fundamental structural niche, and we employ phylogenetic methods to estimate the strength of the association between fundamental and broad-scale realized thermal niche divergence.

## Methods

We measured physiological traits for 304 individuals across 16 species from three islands (Fig. 1; see Supplementary Table 1 for species information). Throughout, we focus primarily on the eight Puerto Rican members of the *cristatellus* group, for two reasons. First, phylogenetic and natural history data indicate that these species diversified *in situ* and form four sister species pairs [46-48], with each pair representing an independent divergence in realized thermal niche based on the degree to which they occupy open versus shaded perches (which throughout we refer to as “warm niche” and “cool niche” species, respectively)[12, 41, 49, 50]. Second, we have detailed measurements of operative thermal environments in Puerto Rico to investigate performance consequences of physiological divergence. Experiments in Puerto Rico were conducted May 20 – June 20 2011, October 8-21 2011, and May 13-28 2012 at the Mata de Plátano field station. On Jamaica, we sampled five of the six extant endemic species, plus the introduced *A. sagrei*. As with the Puerto Rican species, the endemic Jamaican anoles evolved *in situ* from a single colonization event [51]. Experiments in Jamaica occurred from February 25 – March 13 2013 at Green Castle Estates, a privately owned farm and nature preserve in St. Mary Parish. Prior to experiments, lizards were kept individually in plastic cages (18 x 11 x 15 cm) with a wooden dowel perch on an approximately 12L: 12D light schedule. Lizards were watered daily and fed crickets or Phoenix worms every other day. The exception was *A. acutus* from St. Croix, which were brought to Durham, NC and housed following Gunderson and Leal [52].

Heat tolerance (*CT*_max_) was measured following Leal and Gunderson [53], with N = 9-11 individuals/species (Supplementary Table S1). Briefly, lizards (all captured three days prior to measurements) were warmed under a heat lamp while monitoring their body temperatures with a wire thermocouple probe placed inside the cloaca. Lizards were flipped onto their backs at one-degree body temperature intervals starting at 34°C, and *CT*_max_ was recorded as the body temperature at which they lost righting ability. Warming rates were very similar among species, with mean rates ranging from 1.8 to 2.6°C/min (Fig. S1). We modeled variation in log-transformed *CT*_max_ among species using analysis of variance (ANOVA). Planned orthogonal contrasts were applied to test *a priori* predictions about divergence in *CT*_max_ between sister species (in Puerto Rico) and divergence among sympatric, morphologically similar species (Puerto Rico and Jamaica).

Temperature-dependent sprint speeds were measured for thirteen species (N = 9-13 individuals/species, Supplementary Table S1) following Gunderson and Leal [52]. Briefly, we analyzed high-speed video (120 frames/s) of lizards running up a 2 m wooden racetrack set at a 37° angle and marked with a line every 12.5 cm. Lizards were run 2-4 times at each temperature (two runs minimum, additional runs were added if the lizard stopped or jumped off of the track during a trial). Speed for a lizard at a given temperature was taken as the fastest 25 cm interval at that temperature [54, 55]. Puerto Rican lizards were run at five body temperatures in the following randomized order: 32, 22, 27, 17, and 35°C. Conditions were the same for Jamaican lizards, except the 17°C temperature was excluded. All individuals were captured one day prior to the start of trials, and each temperature treatment was administered on a different day over five straight days (four days for Jamaican lizards). No animals were used for both heat tolerance and sprint performance. Target body temperatures were achieved by placing lizards into a calibrated chilling-heating incubator (Cole-Parmer©, Vernon Hills, IL, USA) for 30 min prior to a run [52].

The body temperature of maximum performance (the “optimal temperature,” *T*_opt_) was estimated for each individual [sensu 56]. Thermal performance curves are non-linear [57], and we initially fit two different non-linear models to the sprint data for each individual: a second order polynomial and a modified Gaussian function 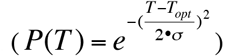) commonly used to fit performance curves for ectotherms [58], including *Anolis* sprinting [52, 59]. The Gaussian model provided relatively poor fits to the data (much higher residual standard errors than the second-order polynomial model (Fig. S2)). Therefore, *T*_opt_ estimates from the second-order polynomial models were used in analyses. *T*_opt_ estimates were constrained such that *T*_opt_ had to occur within the range of experimental temperatures. This is a conservative assumption that likely leads us to underestimate divergence in thermal physiology between species (see Results). *T*_opt_ data were heavily skewed (see Results), so we compared *T*_opt_ among species by assessing overlap in bootstrapped 95% confidence intervals (999 bootstrap replicates using the adjusted bootstrap percentile method due to the skew in our data [60]). Analyses were conducted with the “boot” and “boot.ci” functions, respectively, from the “boot” package in R [61].

We calculated the expected physiological performance of Puerto Rican species under natural warm (open xeric forest) and cool (shaded mesic forest) operative thermal regimes found on Puerto Rico [62]. Operative temperature distributions provide a quantitative description of a thermal environment integrating the physical features of the organism [63], representing the distribution of available body temperatures in a habitat. Operative temperature distributions came from a previous study [52] and were measured with copper lizard models [50] during the peak breeding season (summer) for Puerto Rican anoles [64]. Mean operative temperature in the mesic forest (28.9°C) is significantly cooler than that in the xeric forest (33.4°C) by 4.5°C [52].

We estimated two performance metrics under the different operative environments: 1) relative sprint performance capacity [52, 59, 63], and 2) probability of overheating (the percentage of operative temperature observations over *CT*_max_). For relative performance capacity, we generated species-level thermal performance curves using our sprint speed and *CT*_max_ data. The curves were a combination of two functions fit to the data: one describing performance below *T*_opt_, and the other describing performance from *T*_opt_ to *CT*_max_ [52, 58, 59]. The former function was a second-order polynomial fit to the sprint data, and the latter function took the form: 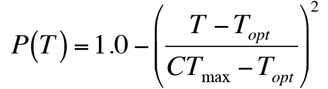 [52, 59]. Curves were scaled to relative performance by setting sprint speed at *T*_opt_ to 1.0 (see Fig. S3 for fitted curves). Performance in each habitat was calculated by applying the curves to operative temperatures [52, 59, 63].

To compare rates of evolution of thermal physiology to ecomorphological traits involved in fundamental structural niche partitioning, we used a phylogenetic comparative approach. Rate of evolution of heat tolerance (*CT*_max_, with expanded species sampling, see Supplementary Table S1) was compared to rates for body size (snout-vent-length, or SVL), head length, femur length, and adhesive toepad width for the 4^th^ hindtoe. We used the maximum clade credibility phylogeny and morphological data of Mahler *et al*. [47] for these analyses. Shape data (head length, femur length, and toepad width) consisted of residuals from phylogenetic regressions of each shape variable on SVL, a standard measure of lizard body size; these residuals were obtained using the phyl.resid function in phytools [65]. We used the approach of Adams [66] to conduct pairwise rate comparisons for all combinations of these five traits. We used likelihood ratio tests to compare a model in which the two traits could evolve at different rates to a model in which the traits were constrained to evolve at a single rate. Rates were estimated using species means of natural log-transformed values for all traits (such that rate variation is represented in terms of relative change in proportion to the trait mean; this is essential for comparison of traits measured using different units [66]). For these analyses, we co-estimated among-trait covariances (similar results were obtained with covariances fixed at zero).

To test for an association between fundamental and broad-scale realized thermal niche estimates, we conducted a phylogenetically-controlled correlation test between *CT_max_* and geo-referenced Worldclim temperature data for each species based on collection localities. We used the first principal component axis from the *Anolis* climate temperature dataset of Algar and Mahler (2015), in which a phylogenetic principal component analysis (PCA) was used to condense the 11 Worldclim variables related to temperature (at 1 km resolution) into PCA axes (Supplementary Table S2). The first PC axis loaded positively for all variables related to temperature (e.g., mean annual temperature, mean maximum temperature; Supplementary Table S2), and can therefore be considered a composite estimate of habitat temperature. Phylogenetically-controlled correlation analyses between *CT_max_* and individual Worldclim variables that loaded strongly with PC1 yielded similar results (Supplementary Figure S4).

## Results

All Puerto Rican sister species pairs have diverged in at least one physiological trait (*T*_opt_, *CT*_max_, or both), and in the predicted direction in all cases (Fig. 2). *T*_opt_ differed between the *A. evermanni/A. stratulus* and *A. cristatellus/A. cooki* species pairs (non-overlapping bootstrapped 95% CIs), but not between the *A. poncensis/A. gundlachi* or *A. pulchellus/A. krugi* species pairs (Fig. 2A). We note that the *T*_opt_ of *A. cooki* and *A. stratulus* were likely underestimated because most individuals had a *T*_opt_ of 35°C solely because that was our maximum test temperature. *CT*_max_ differed among Puerto Rican species (*F*_7,72_ = 42.74; *P* < 0.001), with significant divergence in three of the four species pairs: *A. evermanni/A. stratulus*, *A. poncensis/A. gundlachi*, and *A. pulchellus/A. krugi* (all *P* < 0.001), but not in the *A. cristatellus/A. cooki* species pair (*P* = 0.490; Fig. 2B).

**Figure 2:**
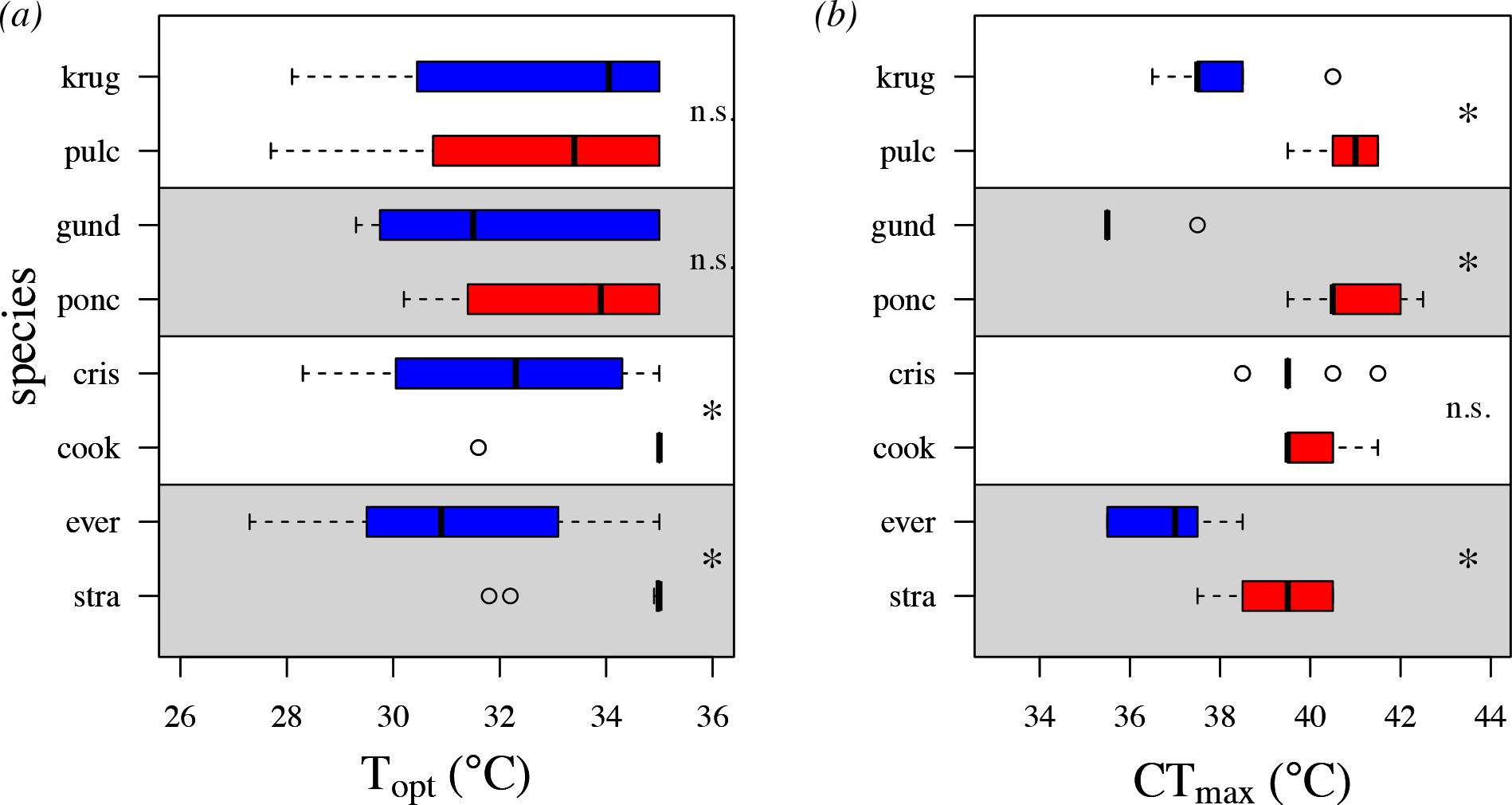
Thermal physiology of four sister-species pairs of Puerto Rican anole. Each species pair is represented in alternating grey and white regions of the plot with warm niche species in red and cool niche species in blue. (A) Optimal sprint performance temperatures (*T*_opt_). (B) Heat tolerance limits (*CT_max_*). “^*^”, significant difference between members of a species pair; N.S., no significant difference between members of a species pair. See Methods for analysis details.

Physiological divergence influenced predicted performance in natural environments. In three of four species pairs, the cool niche species had higher predicted performance than the warm niche species in the shaded operative thermal environment, with mean performance advantages of 8-14% depending on the species pair and time of day (Fig. 3). The lone exception was the *A. pulchellus/A. krugi* species pair, for which both have similar performance estimates (Fig. 3G). Warm niche species tend to have a performance advantage in the open operative environment, particularly during midday hours (from 10:00-13:00; Fig. 3). The *A. cristatellus/A. cooki* species pair is the exception (Fig. 3D). However, *A. cooki* midday performance is likely underestimated due to our probable underestimation of its *T*_opt_ (see above).

**Figure 3:**
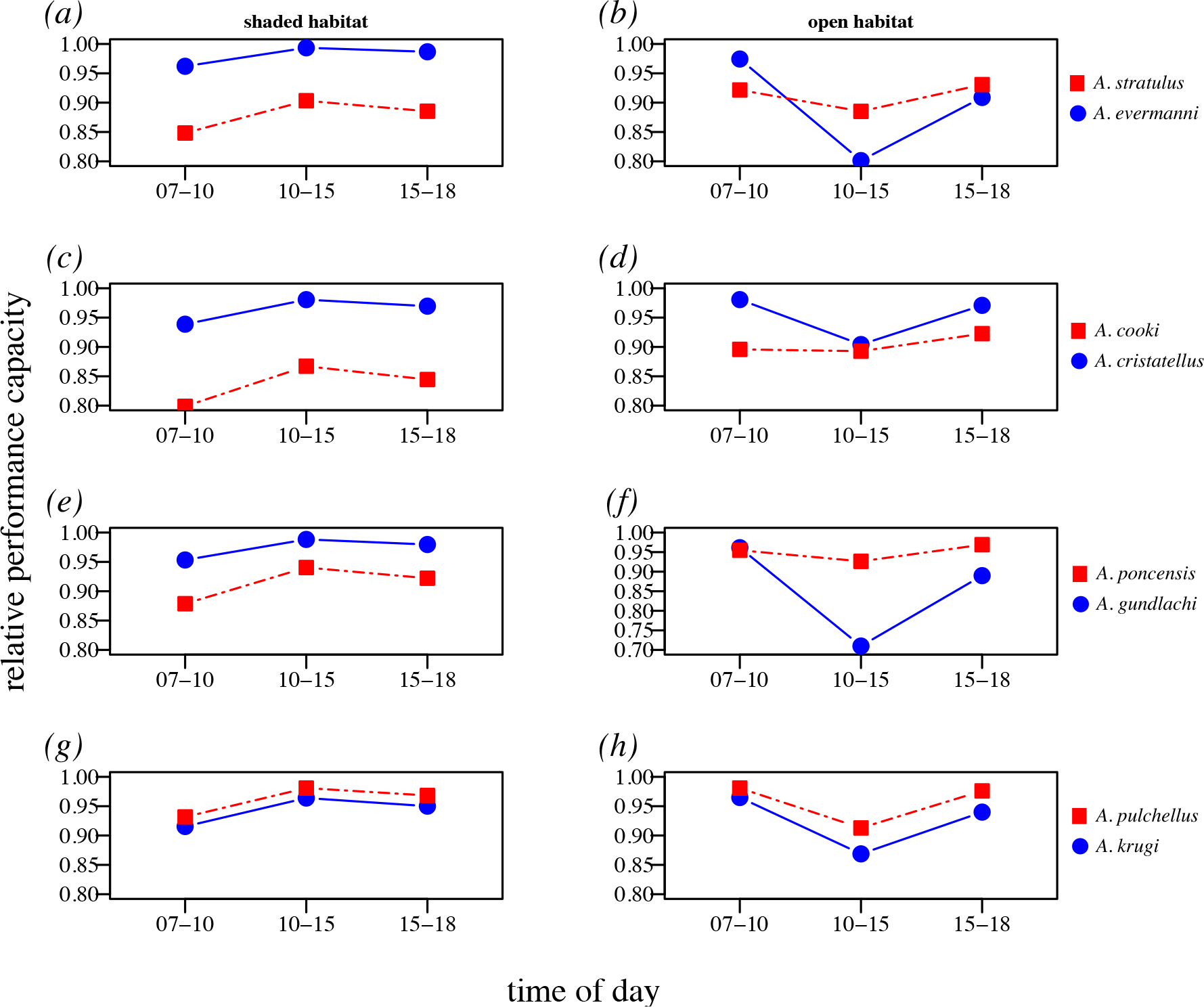
Predicted relative physiological performance of sister-species pairs of Puerto Rican *Anolis* under shaded and open habitat operative thermal environments found on Puerto Rico during morning, midday, and afternoon/evening hours. Warm niche species in red, cool niche species in blue. (A, B) *A. stratulus/A. evermanni*, (C, D) *A. cooki/A. cristatellus*, (E, F) *A. poncensis/A. gundlachi* (G, H) *A. pulchellus/A. krugi*.

No species are under threat of overheating in the shaded operative environment (Fig. 4A, C, E, G). However, all species have overheating risk in the open operative environment (Fig. 4B, D, F, H). For three of the four sister-species pairs (A. *stratulus/A. evermanni*, *A. poncensis/A. gundlachi*, and *A. pulchellus/A. krugi*), the cool niche species has a higher probability of overheating (Fig. 4B, F, H). Difference in overheating probability are particularly acute during midday, when cool niche species have overheating probabilities at least twice that of their warm niche counterparts (G-tests, all *P* < 0.05). The *A. cristatellus/A. cooki* species pair, which had no divergence in *CT*_max_, was the only exception (Fig. 4D).

**Figure 4:**
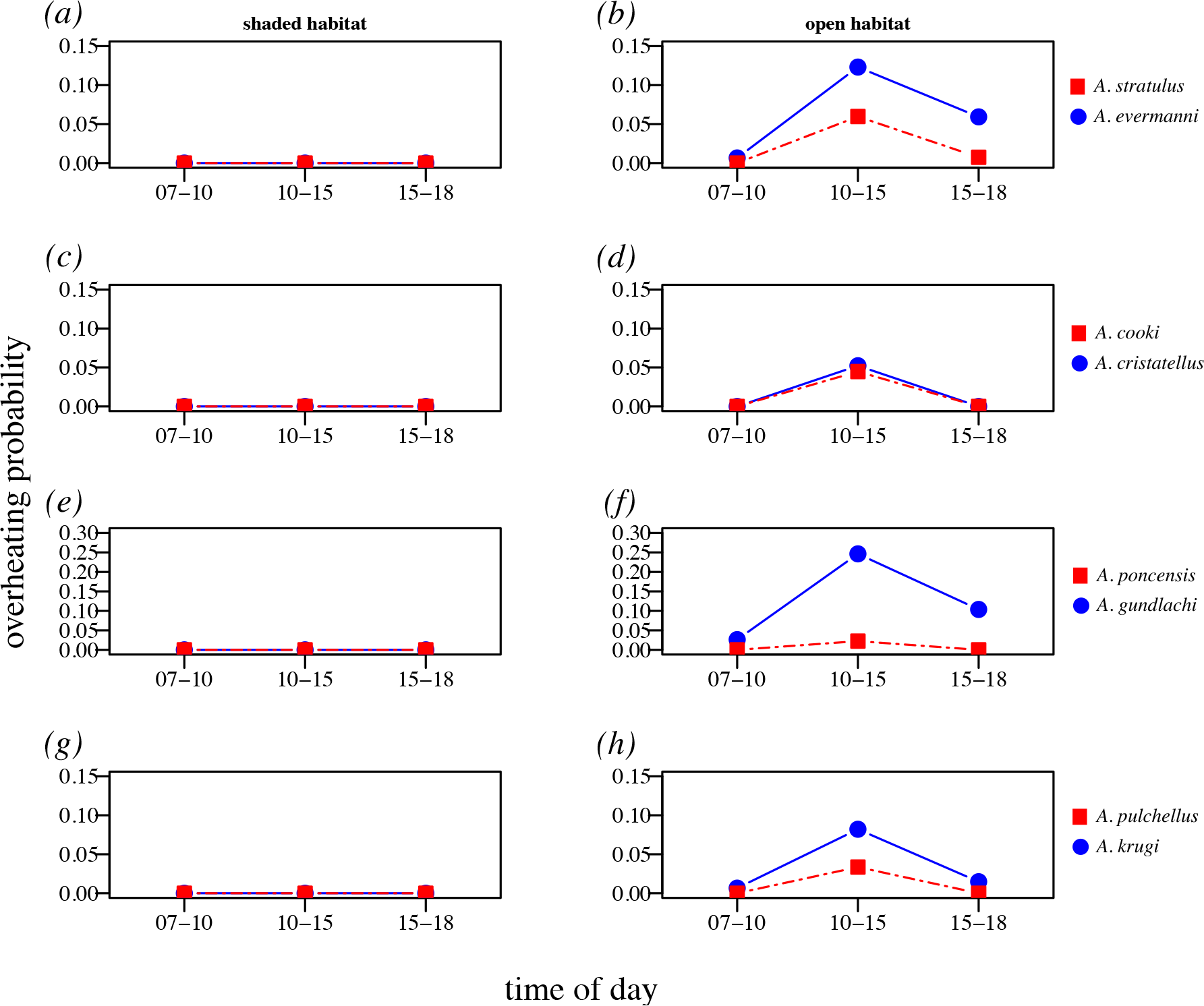
Predicted overheating probability of sister-species pairs of Puerto Rican *Anolis* under shaded and open habitat operative thermal environments found on Puerto Rico during morning, midday, and afternoon/evening hours. Warm niche species in red, cool niche species in blue. (A, B) *A. stratulus*/*A. evermanni*, (C, D) *A. cooki/A. cristatellus* (E, F), *A. poncensis/A. gundlachi*, (G, H) *A. pulchellus/A. krugi*.

Significant physiological divergence is also apparent among Jamaican species. Among species mean *T*_opt_ ranges from 29.5 to 34.6°C (Fig. 5A) while *CT*_max_ ranges from 35.8-41.5°C (Fig. 5B; ANOVA, *F*_336_ = 13.6, *P* < 0.001). Note that *T*_opt_ values of *A. grahami* and *A. opalinus* are likely underestimated due to our upper experimental limit of 35°C (see above).

**Figure 5:**
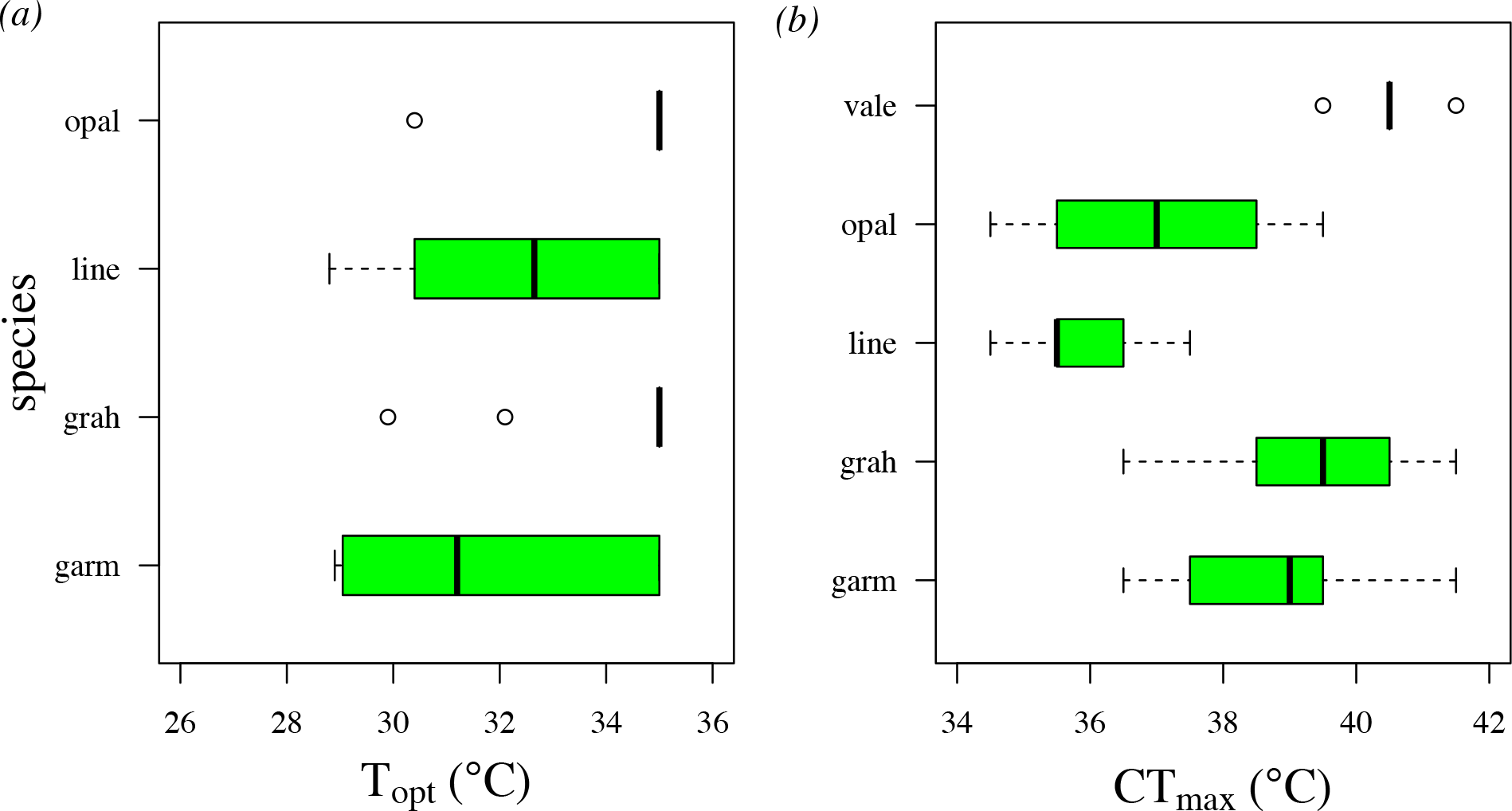
Thermal physiology of endemic Jamaican anoles. (A) Optimal sprinting temperature (*T*_opt_). *Anolis valencienni* does not have a *T*_opt_ estimate because this species could not be induced to run. (B) Heat tolerance (*CT*_max_).

Across Puerto Rico and Jamaica, sympatric species that share the same ecomorphology invariably differ in thermal physiology (Table 1). Thermal tolerance has evolved significantly more slowly than some, but not all, ecomorphological traits (Table 2). *CT*_max_ evolved significantly more slowly than body size (SVL; *P* < 0.001) and relative limb length (femur; *P* = 0.029), but did not differ from head length (*P* = 0.105) or toepad width (*P* = 0.112).

**Table 1.** Physiology differences between co-occurring species that occupy the same or similar structural niches (*i.e.*, share the same or similar ecomorphology). For *CT*_max_, P-values are given based on one-way ANOVA with planned orthogonal contrasts. For *T*_opt_, “^*^” indicates that the bootstrapped 95% confidence intervals did not overlap, “N.S.” indicates no significant difference. “†” indicates that the species is an introduced member of the community.

| co-occurring species |  | ecomorph | physiological trait |  |
| --- | --- | --- | --- | --- |
|  |  |  | CT<sub>max</sub> | T<sub>opt</sub> |
| *A. stratulus* | *A. evermani* | trunk-crown | <0.001 | * |
| *A. cristatellus* | *A. cooki* | trunk-ground | 0.490 | * |
| *A. cristatellus* | *A. gundlachi* | trunk-ground | <0.001 | n.s. |
| *A. krugi* | *A. pulchellus* | grass-bush | <0.001 | n.s. |
| *A. lineatopis* | *A. grahami* | trunk-ground/trunk-crown | <0.001 | n.s. |
| *A. lineatopis* | *A. sagrei*<sup>†</sup> | trunk-ground | <0.001 | n.s. |

**Table 2.** Estimated rates of evolution for heat tolerance (*CT*_max_) and morphological traits related to ecomorphological divergence. Rate estimates (bold) are in the diagonal of the matrix. Upper off-diagonals contain estimates of pairwise evolutionary covariances among traits, lower off-diagonals contain P-values for likelihood ratio test comparisons of 2-rate versus 1-rate models for each trait pair. P-values less than 0.05 indicate support for a 2-rate model over a 1-rate model.

| | $CT_{\max}$ | SVL | head length | femur length | toepad width |
| --- | --- | --- | --- | --- | --- |
| $CT_{\max}$ | **5.80E-05** | -4.90E-05 | -1.60E-05 | -5.50E-05 | -9.90E-06 |
| SVL | <0.001 | **2.30E-03** | 8.90E-06 | -1.80E-05 | 2.70E-05 |
| head length | 0.105 | <0.001 | **2.70E-05** | 1.30E-05 | -6.30E-06 |
| femur length | 0.029 | <0.001 | 0.001 | **1.40E-04** | 2.20E-05 |
| toepad width | 0.112 | <0.001 | 0.003 | 0.832 | **1.30E-04** |

*CT*_max_ was significantly positively correlated with geo-referenced climatic temperatures as represented by temperature principal component axis 1 (*P* = 0.006, r^2^ = 0.45; Fig. 6; see Supplementary Figure S4).

**Figure 6:**
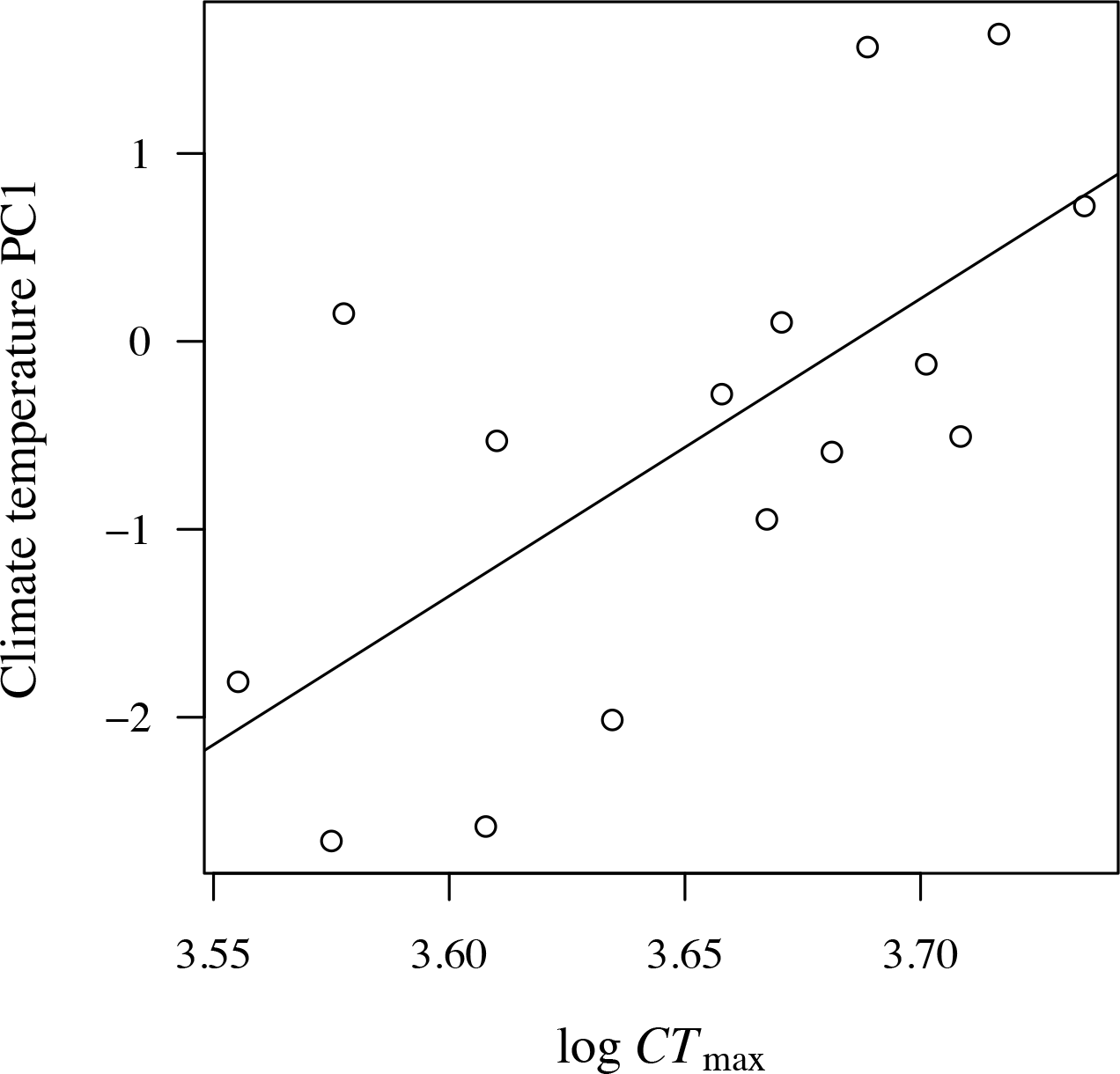
Relationship between *CT*_max_ and climate temperature PC1 from Algar and Mahler (2015).

## Discussion

Abiotic niche evolution has been invoked as an important component of many evolutionary radiations, but the functional consequences of abiotic niche divergence are generally poorly known. We demonstrate that the Puerto Rican *Anolis* adaptive radiation is accompanied by physiological divergence in both heat tolerance and sensitivity to sub-lethal temperature variability. Warm-niche species (determined based on preference for open versus shaded perches) have either higher heat tolerance, higher optimal physiological temperatures, or both, compared to cool-niche species. The anoles of Jamaica also diversified physiologically, with species evolving essentially the full range of heat tolerances and optimal temperatures observed in their Puerto Rican counterparts (Fig. 5). These results highlight the possibility that physiological adaptation can be pervasive within evolutionary radiations, including those for which morphological divergence is widespread.

Repeated divergence in physiological traits under similar conditions is itself evidence of adaptation [67]. However, we provide more direct evidence of adaptation by estimating physiological performance of species in warm and cool operative environments on Puerto Rico. Cool niche species have higher predicted performance in shaded operative environments than their warm niche sister species in three of the four species pairs (Fig. 3), with the lone exception a species pair for which there was no divergence in *T*_opt_. The pattern was less pronounced in the open operative environment, with warm niche species having a sizable performance advantage in only two species pairs (Fig. 3). However, the performance advantage for warm niche species becomes apparent when considering overheating risk. During midday hours in the warmer open environment, warm niche species in three of the four species pairs had less than half the overheating risk of cool niche species (Fig. 4). The sole exception was the *A. cristatellus/A. cooki* species pair, which exhibited no divergence in thermal tolerance (Fig. 2B). One implication of these results is that what appears to be modest physiological evolution (*CT*_max_ and *T*_opt_ divergence of 2-4°C) can still have significant performance consequences.

The high community-level species diversity associated with *Anolis* and other adaptive radiations is facilitated by fine-scale resource partitioning [3]. Sympatric anoles are known to behaviorally partition thermal microhabitats [12, 13], and our findings indicate that this is facilitated by physiological divergence. When co-occurring anole species share the same fundamental structural niche (*i.e*., ecomorphological microhabitat specialization), they invariably differ in thermal physiology (Table 1).

This pattern is also maintained for introduced species: in Jamaica, the endemic trunk-ground species *A. lineatopus* is found in sympatry with the introduced trunk-ground species *A. sagrei*, and these species differ significantly in thermal tolerance (Table 1). These data highlight that, even in radiations for which morphological divergence is important in facilitating coexistence, physiological divergence can also play an important role, providing novel axes of diversification [68].

We found that thermal tolerance evolved more slowly than some ecomorphological traits (*i.e.*, SVL and femur length), and evolved at similar rates to others (*i.e.*, head length and toe-pad width; Table 2). In general, divergence along the fundamental thermal niche axis appears to have occurred more slowly than divergence along the fundamental structural niche axis, at least within the Puerto Rican and Jamaican anole radiations. This finding is inconsistent with a recent analysis that found that thermal niche, estimated as field body temperature, evolves more quickly than morphology in Caribbean anoles [43]. Discrepancy between these results may be due to the different traits analyzed and/or the different species included, or the fact that that study estimated rates of absolute, rather than proportional, trait change (Fig. 4). Regardless, the observation that physiology evolves slowly, with repeated divergence of relatively small magnitude, is consistent with our above suggestion that large changes in physiology are not necessary for divergence to be ecologically important. These results also indicate that caution should be exhibited when inferring the importance of traits based on comparisons of evolutionary rate estimates in the absence of data on how phenotypic changes map to performance change under natural conditions.

We found that fundamental and broad-scale realized thermal niche estimates were correlated, as *CT*_max_ was significantly positively correlated with geo-referenced temperature data across species (Fig. 6). Therefore, the realized thermal niches estimated in correlative ENM studies can capture some of the underlying variability in the fundamental thermal niche. Nonetheless, the majority of variability in the data (∼55%) is left unexplained, and there are clear cases where realized and fundamental niche estimates diverge. For example, *A. lineatopus* and *A. gundlachi* have very similar *CT*_max_ (mean *CT*_max_ = 36.3°C and 36.2°C, respectively), but occur in very different realized thermal niches (temperature PC1 = 0.14 and - 2.66, respectively). Conversely, *A. opalinus* and *A. pulchellus* occur in very similar realized niches (temperature PC1 = −0.53 and −0.51, respectively) but have very different *CT*_max_ (mean *CT*_max_ = 37.5°C and 41.3°C, respectively). These results underscore the fact that broad-scale climatic conditions are not necessarily reflective of the underlying thermal biology of the organism being considered. Much of this discrepancy is likely due to behavioral thermoregulation, which allows taxa in the same geographic location to experience very different body temperatures and those in different geographic locations to experience very similar body temperatures [35, 69].

We have shown that fundamental thermal niche divergence is important in the Puerto Rican and Jamaican *Anolis* adaptive radiations. However, the importance of such divergence is likely not restricted to these islands. For example, species of *cybotes* group trunk-ground anoles in Hispaniola have diverged in thermal tolerance along an elevational gradient [45, 70], and variation in thermal physiology also occurs among non-Antillean anoles [71, 72]. Water loss rates also differ among anole species and populations [73-79]. In this context, our results suggest that physiological divergence may be as important as morphological divergence for diversification and coexistence in this classic radiation.

Achieving a mechanistic understanding of how evolutionary and ecological processes interact to promote the production and maintenance of biodiversity is a long-standing goal in evolutionary ecology. Given the reality of ongoing climate change, the need to understand these processes has become immediate, particularly with respect to temperature-dependent traits [80]. By focusing on physiological traits that link temperature to performance, we demonstrate that the evolution of thermal physiology can facilitate adaptive radiation by contributing to *in situ* performance tradeoffs and species co-existence. While our trait- and performance-based analyses would not be possible using the type of broad-scale climatic data used in ENMs studies alone, we find that climate-scale realized niche features correlate with fundamental niche features. Nonetheless, climatic data provide imperfect signal with respect to the evolution of functional traits, likely due to the ability of species to behaviorally augment their effective thermal environments. We suggest that deeper insights about the contribution of climatic niche divergence to evolutionary radiations will emerge when correlative niche data are used in tandem with experimental physiological studies within an integrative research program.

## Acknowledgements

We thank J. Losos and two anonymous reviewers for helpful comments on this manuscript, A. Algar for help with data, D. Steinberg for field help and the El Verde and Mata de Plátano field stations, Ridge to Reef farm, and Green Castle Estates for logistical support. Permits were provided by the Puerto Rican Departmento de Recursos Naturales y Ambientales, the St. Croix Division of Fish and Wildlife, and the Jamaican National Environment and Planning Agency.

**Funding.** This research was funded by NSF DDIG #1110570 to A.R.G.

**Ethics.** This research was conducted with the approval of the Duke University Animal Care and Use Committee permit #A106-10-04.

**Author Contributions**. A.R.G. and M.L. designed the study. A.R.G. collected the physiological data and conducted all statistical analyses that were not phylogenetically controlled. L.D.M. conducted all phylogenetic comparative analyses. A.R.G. led the writing with contributions from M.L. and L.D.M.

**Competing interests**. We have no competing interests.

**Data accessibility**. Physiological data are deposited in Dryad (DOI: https://doi.org/10.5061/dryad.688jj72)

